# Diet-dependent *Natriuretic Peptide Receptor C* expression in adipose tissue is mediated by PPARγ via long-range distal enhancers

**DOI:** 10.1101/2021.02.09.430332

**Authors:** Fubiao Shi, Zoltan Simandi, Laszlo Nagy, Sheila Collins

## Abstract

In addition to their established role to maintain blood pressure and fluid volume, the cardiac natriuretic peptides (NPs) can stimulate adipocyte lipolysis and control the brown fat gene program of nonshivering thermogenesis. The NP “clearance” receptor C (NPRC) functions to clear NPs from the circulation via peptide internalization and degradation and thus is an important regulator of NP signaling and adipocyte metabolism. It is well appreciated that the *Nprc* gene is highly expressed in adipose tissue and is dynamically regulated with nutrition and environmental changes. However, the molecular basis for how *Nprc* gene expression is regulated is still poorly understood. Here we identified Peroxisome Proliferator-Activated Receptor gamma (PPARγ) as a transcriptional regulator of *Nprc* expression in mouse adipocytes. During 3T3-L1 adipocyte differentiation, levels of *Nprc* expression increase in parallel with PPARγ induction. Rosiglitazone, a classic PPARγ agonist, increases, while siRNA knockdown of PPARγ reduces, *Nprc* expression in 3T3-L1 adipocytes. We demonstrate that PPARγ controls *Nprc* gene expression in adipocytes through its long-range distal enhancers. Furthermore, the induction of *Nprc* expression in adipose tissue during high-fat diet feeding is associated with increased PPARγ enhancer activity. Our findings define PPARγ as a mediator of adipocyte *Nprc* gene expression and establish a new connection between PPARγ and the control of adipocyte NP signaling in obesity.

## Introduction

The cardiac natriuretic peptides (NP), including atrial NP (ANP) and B-type NP (BNP), were first discovered as factors in atrial extracts that evoked a strong decrease in blood pressure when injected into rodents (1). They are now known to be secreted from atrial cardiomyocytes in response to increases in blood volume and cardiac wall stress. However, in conditions of persistent high blood pressure, some reports suggest BNP can also be produced in the ventricle (2). Thus the most well established physiological function of NPs is to maintain blood pressure and fluid volume (3). More recently, NPs have also been shown to stimulate adipocyte lipolysis to liberate fatty acids from adipose tissue with a potency comparable to the catecholamines (4,5). This places the heart in a position of being both a consumer of fatty acids as well as a regulator of their release from adipose tissue. Furthermore, work from our lab has demonstrated that NPs could also promote the thermogenic program of brown adipocytes by increasing the gene expression programs of mitochondrial biogenesis, fatty acid oxidation and the key thermogenic protein uncoupling protein-1 (UCP1) (6).

The physiological effect of the NPs is mediated by the cyclic guanosine monophosphate (cGMP) - protein kinase G (PKG) signaling cascade in target cells. There are two receptors for the cardiac NPs: NP receptor A (NPRA) and NP receptor C (NPRC). Binding of NPs to NPRA activates its intracellular guanylyl cyclase activity, increases cGMP production and triggers the PKG-dependent signaling cascade (7). In contrast, binding of NPs to NPRC, which lack an intracellular guanylyl cyclase domain, results in the internalization and degradation of the peptides (7). NPRC-mediated NP degradation is an important mechanism to modulate the available pool of NPs for target cell activation.

NPRC appears to be an important regulator of adipose tissue NP signaling and energy metabolism. In humans, increases in circulating NPs are associated with weight loss, while obese human subjects across ethnic groups with metabolic syndrome often show reduced circulating NPs and biological efficacy (e.g., elevated blood pressure) (8-11). This biological response to NPs depends on the relative amount of the guanylyl cyclase receptor NPRA to the ‘clearance’ receptor NPRC. In mice it was shown that genetic deletion of NPRC did not change the plasma levels of ANP and BNP, but their circulating half-life was substantially increased (12). In adipocytes, the levels of NPRC are dynamically regulated in response to nutritional and hormonal status. In obese humans and rodent models, the levels of NPRC are elevated in adipose tissue (9,13-18). This is of physiological significance because it has been proposed that greater removal of NPs from circulation by NPRC in adipose tissue could explain what has been referred to as a ‘natriuretic handicap’ linking obesity and hypertension (9-11). We previously reported that deletion of *Nprc* specifically in adipose tissue increases NP signaling and protects against diet-induced obesity and insulin resistance (16). As a result, the level of NPRC in adipose tissues is crucial for NP action. In spite of this important biology, there is little known about how the *Nprc* gene is regulated. To tackle this problem, we used *in vitro* adipocyte culture and diet-induced obesity mouse models to investigate the mechanisms by which *Nprc* gene expression is regulated in adipocytes and adipose tissues. Here we describe our results which identified Peroxisome Proliferator-Activated Receptor gamma (PPARγ) as a transcriptional regulator of *Nprc* expression and showed that the PPARγ enhancer activity in the *Nprc* gene is related to *Nprc* mRNA induction in response to high-fat diet feeding. Although there are probably additional factors to control *Nprc* gene expression, our data provide the first insights on *Nprc* transcriptional regulation and establish a new role for PPARγ to control adipocyte NP signaling and metabolism during obesity.

## Results

### Regulation of *Nprc* expression in adipocytes by PPARγ

In efforts to understand how *Nprc* expression is regulated in adipocytes, we first examined the expression of *Nprc* over the time course of 3T3-L1 adipocyte differentiation. As shown in Fig. 1A, *Nprc* mRNA levels progressively increased throughout the differentiation process. Meanwhile, the expression of *Pparγ*, the master regulator of adipocyte differentiation (19), and *Fatty acid-binding protein 4* (*Fabp4*), a well-characterized PPARγ target gene (20,21), also increased over the course of differentiation (Fig. 1B, 1C). In line with this, although *Nprc* protein was not detectable in the preadipocytes, it increased along with the differentiation process, in parallel to the protein levels of PPARγ and its targets FABP4 and Adiponectin (ADIPOQ) (22) (Fig. 1D). When fully differentiated, adipocytes treated with rosiglitazone, a potent PPARγ agonist, for 6 hours showed a further increase in *Nprc* transcript levels, in a similar fashion to *Fabp4* (Fig. 2A). In contrast, siRNA knockdown of *Pparγ* resulted in a significant reduction of *Nprc* levels compared to the scramble siRNA control (Fig. 2B). Similarly, NPRC protein was also significantly reduced with PPARγ knockdown as expected, like the other two PPARγ targets FABP4 and ADIPOQ (Fig. 2C). These data suggested that PPARγ controls *Nprc* expression in adipocytes. We further confirmed this observation in NIH-3T3 fibroblasts that stably express PPARγ (NIH-PPARγ) (23,24). In NIH-PPARγ cells, *Nprc* mRNA increased in response to rosiglitazone treatment as expected, though to a lesser extent than *Fabp4*. In contrast, in the parental NIH-3T3 cells that lack PPARγ, *Nprc* as well as *Fabp4* transcripts did not increase after rosiglitazone treatment (Fig. 2D). These data further support the conclusion that expression of *Nprc* is regulated by PPARγ.

**Figure 1.**
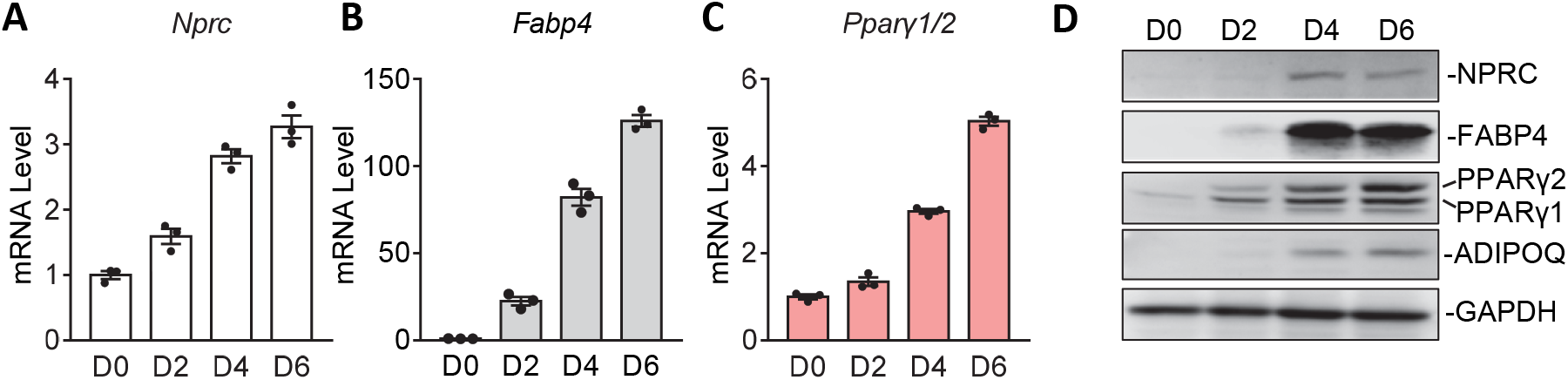
*Nprc* mRNA expression during 3T3-L1 adipocyte differentiation. (**A-C**) mRNA levels of *Nprc* (A), *Fabp4* (B) and *Pparγ½*(C) during 3T3-L1 adipocyte differentiation. Data were normalized with 36B4. (**D**) Protein levels of NPRC, FABP4, PPARγ½and ADIPOQ during 3T3-L1 adipocyte differentiation (Day 0, 2, 4 and 6).

**Figure 2.**
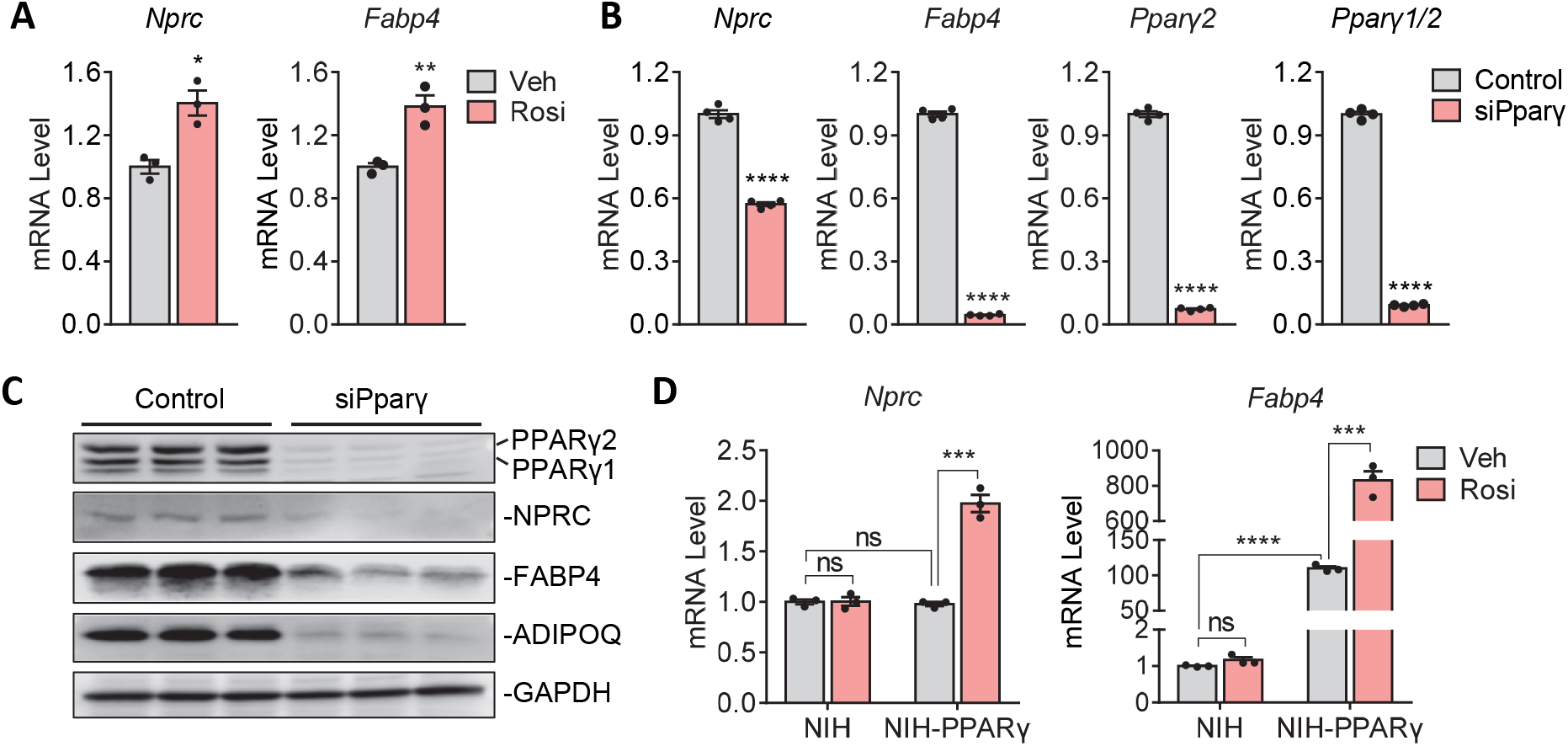
Modulation of *Nprc* expression by rosiglitazone and PPARγ. mRNA levels of *Nprc* and *Fabp4* in 3T3-L1 adipocytes after treatment with 1μM rosiglitazone (Rosi) or vehicle (Veh) for 6 hours. (**B**) mRNA levels of *Nprc, Fabp4* and *Pparγ½*mRNA in 3T3-L1 adipocytes after siRNA knock down of *Pparγ* (siPparγ). (**C**) Protein levels of PPARγ1/2, NPRC, FABP4 and ADIPOQ in 3T3-L1 adipocytes after siRNA knock down of *Pparγ* (siPparγ). (**D**) mRNA levels of *Nprc* and *Fabp4* in NIH-3T3 cells (NIH) and NIH-3T3 cells stably expressing PPARγ (NIH-PPARγ) after treatment with 1μM rosiglitazone (Rosi) or vehicle (Veh) for 6 hours. QPCR data were normalized with 36B4. Student’s t-test, *p<0.05, **p<0.01, ***p<0.001, ****p<0.0001, ns: not significant.

### Interaction of distal PPARγ enhancers with *Nprc* promoter in adipocytes

To determine the mechanism by which PPARγ regulates *Nprc* expression, we next searched for PPARγ binding sites in the mouse *Nprc* locus by analyzing a previously published PPARγ ChIP-seq dataset from 3T3-L1 adipocytes and mouse subcutaneous inguinal white adipose tissue (iWAT) (25). As shown in Fig. 3A, there were several PPARγ binding sites in the regions -9kb, -44kb, -49kb, -54kb, -58kb, -62kb and - 71kb upstream of the *Nprc* transcription start site. These PPARγ binding peaks were well harmonized between 3T3-L1 adipocytes and the iWAT depot. It was also noted that the PPARγ binding activity was higher in the iWAT of mice fed with high-fat diet, and further increased with rosiglitazone administration (Fig. 3A). These data suggest that the control of *Nprc* expression by PPARγ could be mediated by long-range enhancers.

**Figure 3.**
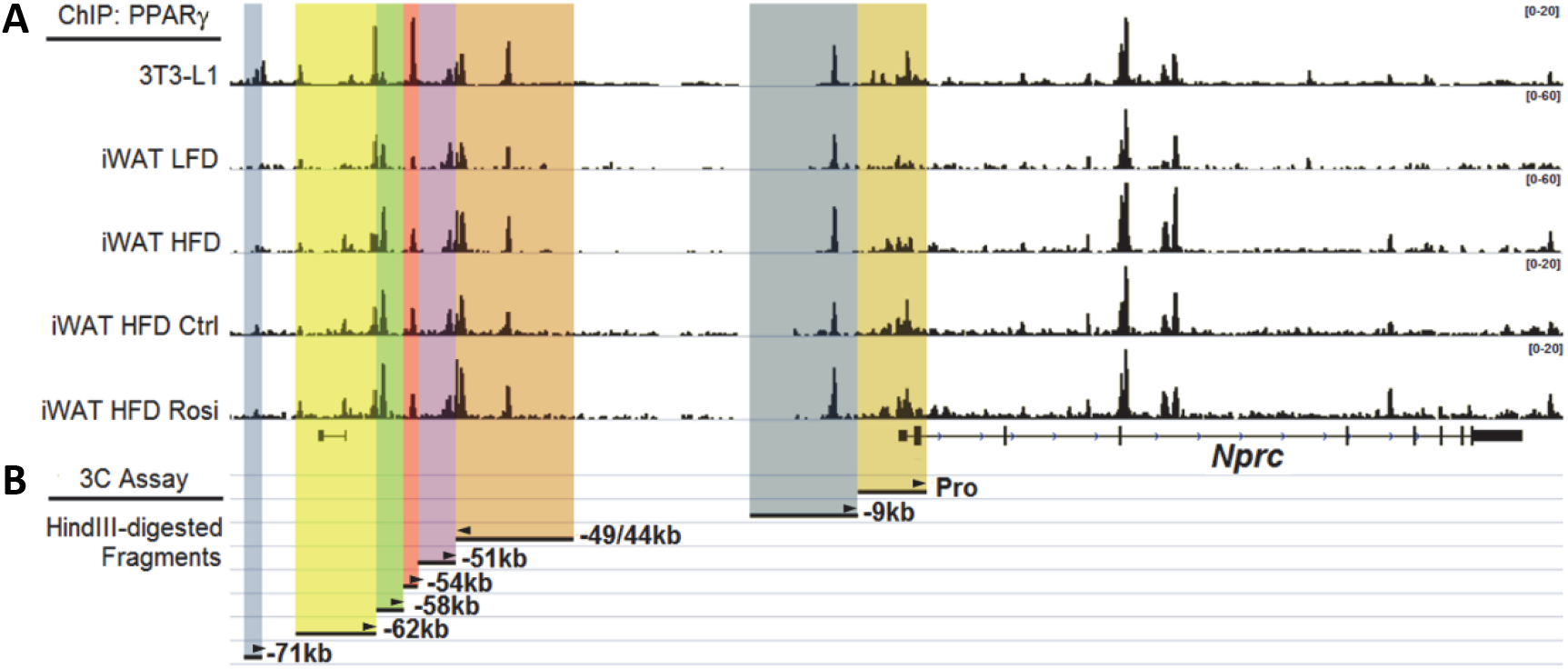
ChIP-seq identified PPARγ enhancers in the upstream distal region of *Nprc*. (**A**) PPARγ binding sites in the upstream distal region of the *Nprc* promoter were identified by ChIP-seq in 3T3-L1 adipocytes and inguinal white adipose tssue (iWAT) from mice fed with low-fat diet (LFD), high-fat diet (HFD), HFD plus vehicle (Ctrl), and HFD plus rosiglitazone (Rosi) (Soccio et al., JCI, 2017). (**B**) HindIII-digested restriction fragments containing *Nprc* promoter (Pro) and the PPARγ binding sites (−9kb, -44/49kb, -51kb, -54kb, -58kb, 62kb and -71kb) are illustrated for the chromosomal conformation capture (3C) analysis in Figure 4. The arrowheads indicate the primers used for 3C PCR in each fragment.

To determine whether these PPARγ binding sites are directly involved in the regulation of *Nprc* expression, we utilized chromosomal conformation capture (3C) assays to examine the interaction of these enhancers with the proximal *Nprc* promoter (Fig. 3B, 4A). Among those HindIII-digested restriction fragments that contained PPARγ enhancers, only the -9kb, -44/49kb and -54kb fragments could form chimeric ligation products with the *Nprc* promoter fragment (Fig. 4B), suggesting that the distal enhancers in these three fragments could interact with the *Nprc* promoter by forming a looping structure, which is essential for the function of distal enhancer elements (26).

**Figure 4.**
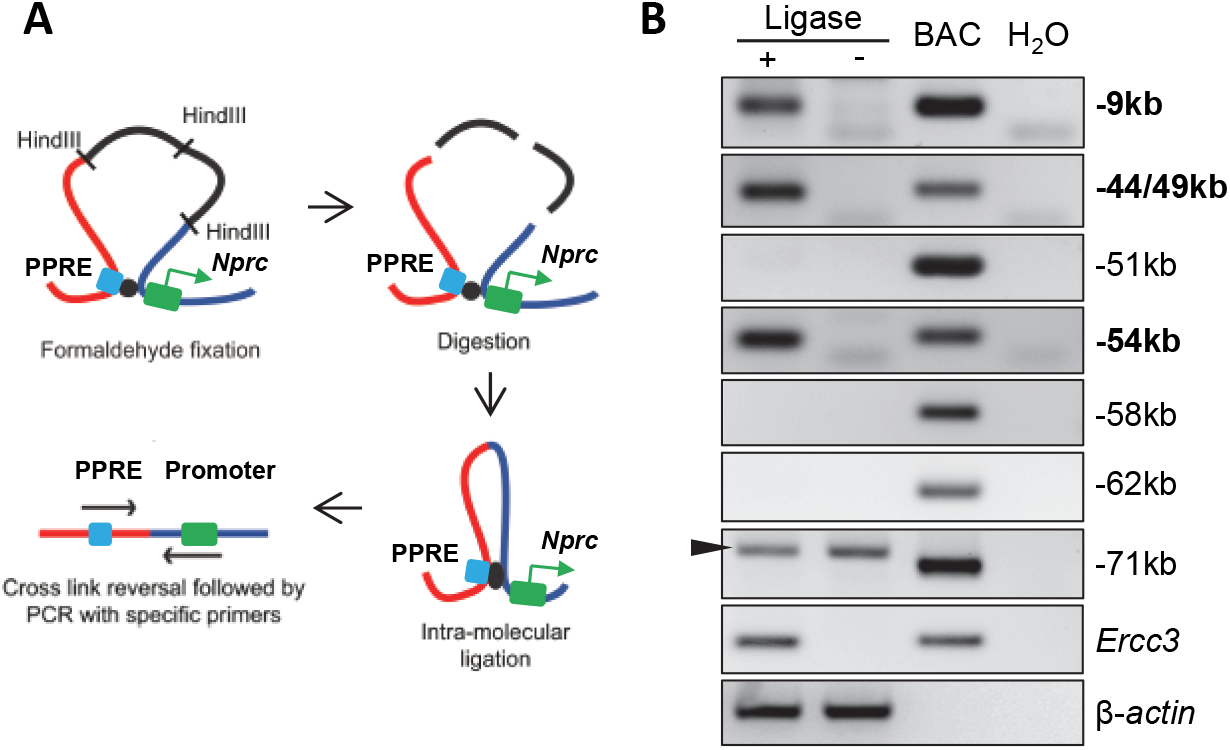
Interaction of PPARγ-binding fragments with *Nprc* promoter. (**A**) Procedure of chromosomal conformation analysis (3C) (modified from Cope and Fraser, 2009). PPRE: PPARγ response elements. (**B**) 3C analysis with C3H10T_½_adipocytes to determine the interaction between *Nprc* promoter and PPARγ-binding sites within each HindIII-restriction fragment (−9kb, -44/49kb, -54kb, -58kb, -62kb and -71kb, also see Figure 3B for details). *Ercc3* was positive control for constitutive enhancer interaction. β-*actin* was DNA input control. Ligase+, HindIII-digested chromatin with intramolecular religation. Ligase-, HindIII-digested chromatin without intramolecular religation. BAC, bacterial artificial chromosome DNA covering *Nprc* (RP23-305L10) and *Ercc3* (RP23-148C24) genes was digested with HindIII and randomly religated as positive control. Arrowhead indicates non-specific amplification products.

### Functionality of *Nprc* promoter and the distal PPARγ enhancers

To determine the functionality of the *Nprc* promoter and these distal enhancers, we next cloned a 2.2kb *Nprc* promoter fragment and the PPARγ enhancer fragments (as showed in Fig. 5A) into a luciferase reporter system to test their reporter activity in response to rosiglitazone treatment. The *Nprc*-2.2k promoter did not show any luciferase activity in NIH-PPARγ cells in response to rosiglitazone (Fig. 5B). The *TK* promoter and the *Angptl4* enhancer were used as negative and positive controls, respectively, as described in a previous report (27). Among all the distal enhancer fragments tested, the -49kb, -54kb and -62kb PPARγ enhancer fragments showed significant increases in luciferase activities in response to rosiglitazone (Fig. 5C). Since the -62kb fragment did not interact with *Nprc* promoter (Fig. 4B), it may be that it participates in the regulation of a gene other than *Nprc*. Therefore, our further analysis focused on the - 49kb and -54kb fragments. More detailed sequence analysis showed that there were three putative PPARγ response element (PPRE) motifs within the -49kb fragment and one in the -54kb fragment (Fig. 5A). These four PPREs were further cloned into the luciferase reporter vector. Both the -49kb-P2 and -54kb-P4 elements showed increased luciferase activity with rosiglitazone treatment, illustrating that the -49kb-P2 and -54kb-P4 are functional PPARγ binding sites, although the other candidate sites in the -49kb region may function only in the context of the intact chromatin. Although the fold induction of the individual PPRE is modest, the agonist effect is very specific and reproducible, and these elements might work together within the context of intact chromatin in a cooperative or synergistic manner. Note that in the parental NIH-3T3 cells, which do no express PPARγ, none of the enhancer and PPRE fragments, including the positive control *Angptl4* enhancer, showed any luciferase activity after rosiglitazone. This is consistent with a specific role for PPARγ in NIH-PPARγ cells. Taken together, these data suggest that PPARγ controls *Nprc* gene expression through these long-range distal enhancers.

**Figure 5.**
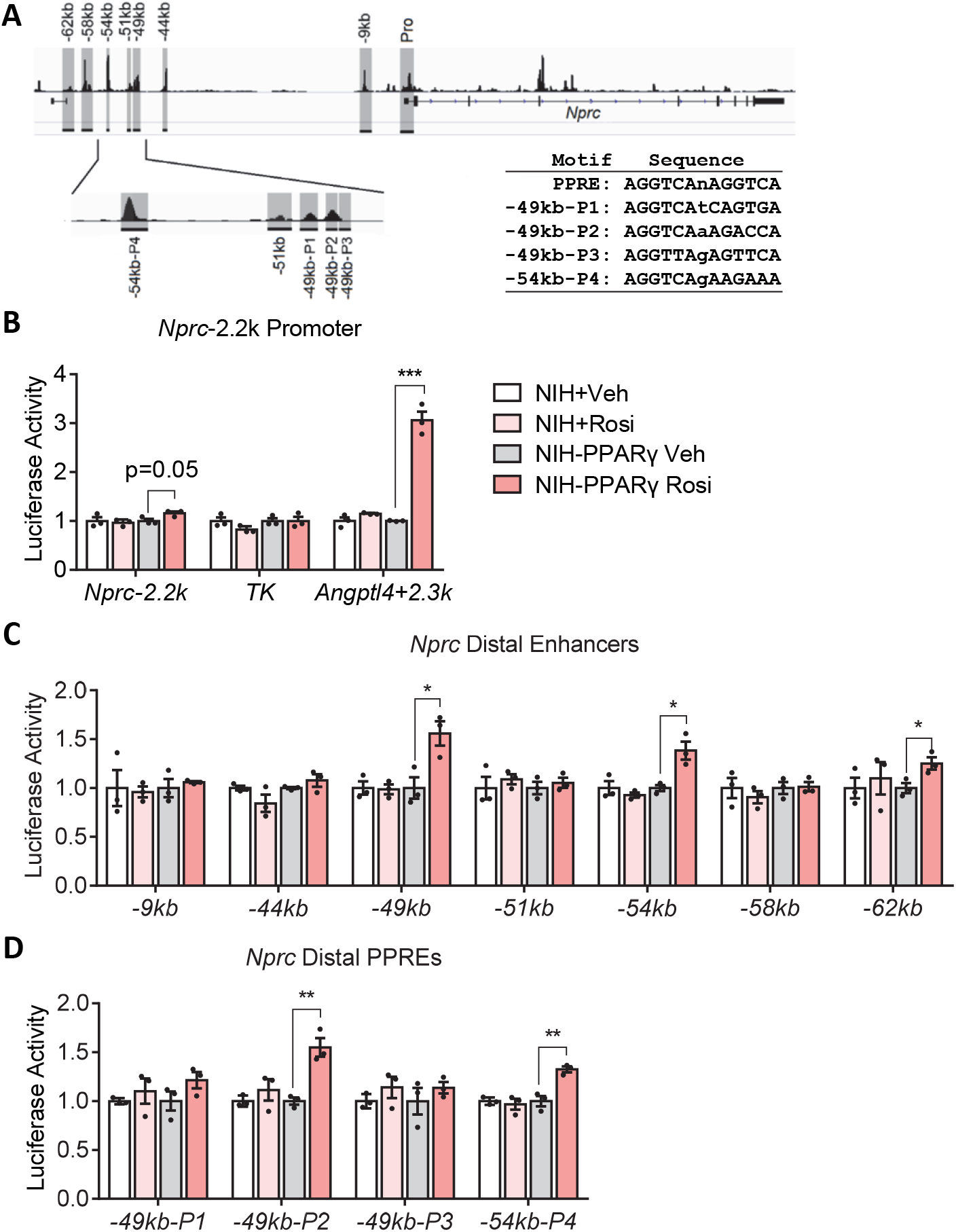
Rosiglitazone increases *Nprc* distal enhancer activity but not proximal promoter activity alone. (**A**) *Nprc* promoter (Pro), distal PPARγ enhancers (−9kb, -44kb, 49kb, -51kb, -54kb, -58kb, 62kb) and the four PPARγ response elements (PPRE, P1, P2, P3 and P4) are shown in gray and were cloned for luciferase reporter analysis. The sequences of the consensus PPRE motif and the four *Nprc* PPREs are listed as insert. (**B**) Luciferase activity of *Nprc* promoter (−2233bp to +1bp, *mNprc-2*.*2k*). *TK* promoter alone (*TK*) and *TK* promoter with a PPARγ enhancer from the *Angptl4* gene (*Angptl4+2*.*3k*) were used as the negative and positive controls, respectively. (**C**) Luciferase activity of the *Nprc* distal enhancer fragments (−9kb, -44kb, -49kb, -51kb, -54kb, -58kb and -62kb). (**D**) Enhancer activity of the *Nprc* distal PPREs in the -49kb fragment (P1-P3) and -54kb fragment (P4). For reporter assays, NIH-3T3 (NIH) and NIH-3T3 stably expressing PPARγ (NIH-PPARγ) cells were transfected with reporter plasmids and treated with vehicle (Veh) or 1μM rosiglitazone (Rosi) for 48 hours. Luciferase activity was normalized to protein concentrations. Data were representative of at least three independent experiments. Student t-test, *p<0.05, **p<0.01, ***p<0.001.

### High-fat diet-induced *Nprc* gene expression in adipose tissue is associated with increased PPARγ enhancer activity

We and others have shown that *Nprc* expression is increased in rodent adipose tissue in response to high-fat diet (HFD) feeding (16-18). Coupled with the observation that there is more PPARγ binding to these distal enhancers in the ChIP-Seq data (Fig 3A; ref 25), we next questioned whether the increase of *Nprc* expression in HFD-fed mice is mediated by PPARγ. To test this hypothesis, we examined the -49kb and - 54kb PPARγ enhancer activities in adipose tissues by measuring their enhancer RNA (eRNA) synthesis via QPCR (28). Interestingly, the eRNA level of -49k PPARγ enhancer was significantly increased in the brown adipose tissue (BAT) and iWAT of HFD-fed mice, in parallel to the changes in *Nprc* gene expression itself (Fig. 6A). The increase of -54k PPARγ eRNAs was also significant in iWAT but only marginally so in the BAT (Fig. 6A). The level of *Fabp4* mRNA was slightly decreased in BAT, but its -5.4kb PPARγ eRNA was not altered with HFD feeding (Fig. 6B), suggested a tissue and locus-specific effect of PPARγ action. The level of *Npra* mRNA was modestly increased in BAT, and unchanged in iWAT with HFD (Fig. 6C). In sum, our results lead us to conclude that obesity increases *Nprc* expression in the adipose tissue through a PPARγ-dependent mechanism, the effect of which is the suppression of NP signaling, decreased energy expenditure and impaired glucose homeostasis (Fig. 7).

**Figure 6.**
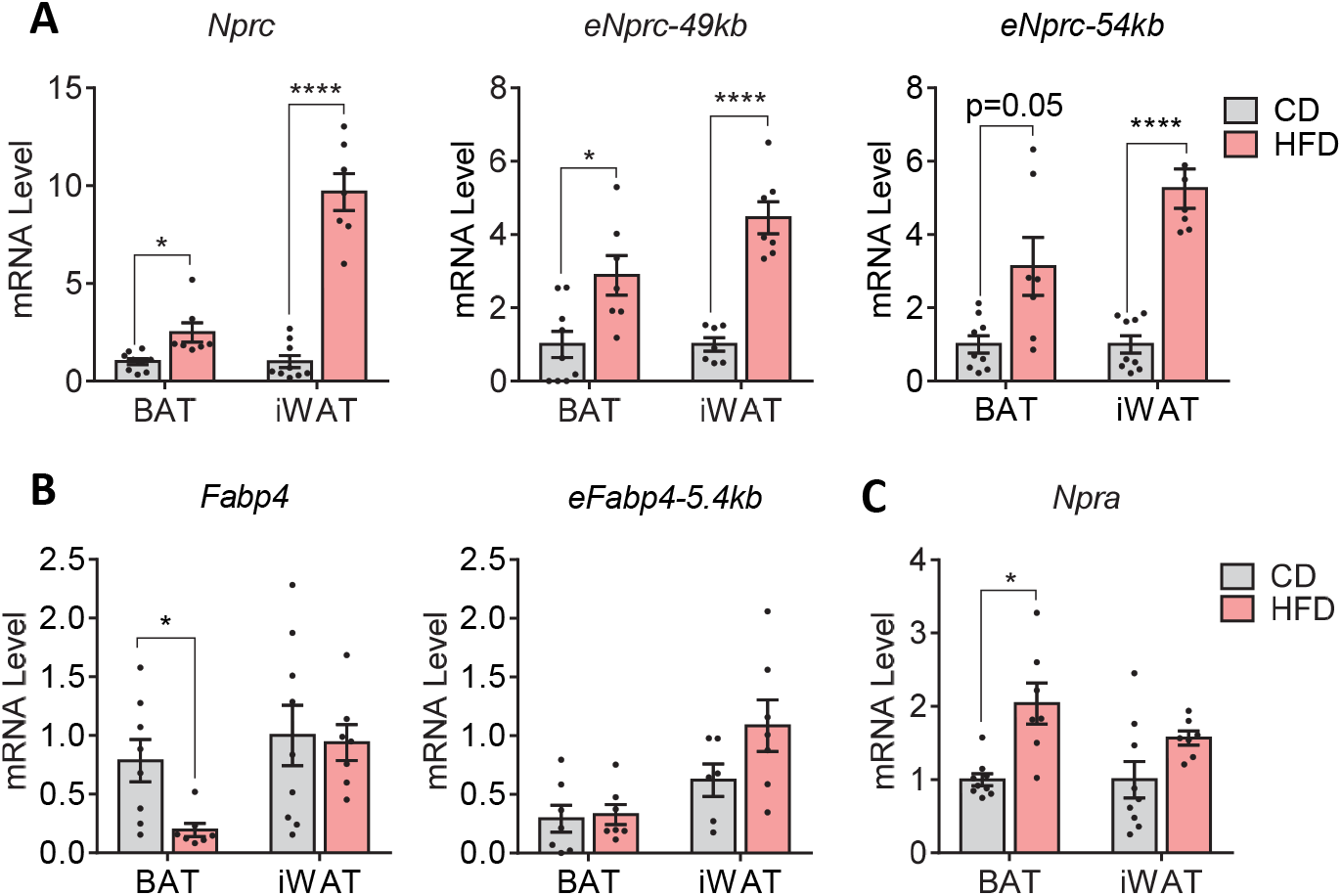
High-fat diet induces *Nprc* expression and PPARγ enhancer activity in adipose tissue. (**A**) Levels of *Nprc* mRNA and the -49kb, -54kb PPARγ enhancer RNAs (*eNprc*) in brown adipose tissue (BAT) and inguinal white adipose tissue (iWAT) of wild-type mice fed with control diet (CD) or high-fat diet (HFD). (**B**) Levels of *Fabp4* and the -5.4kb PPARγ enhancer RNA (*eFabp4*) in BAT and iWAT of mice fed with CD and HFD. (**C**) Levels of *Npra* mRNA in the BAT and iWAT of mice fed with CD and HFD. Student t-test, *p<0.05, ****p<0.0001, ns: not significant.

**Figure 7.**
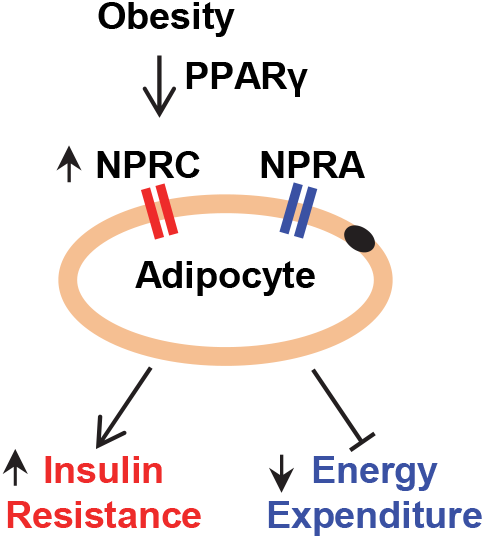
Obesity induces NPRC expression in adipocytes. Obesity induces NPRC expression in adipocytes potentially though a PPARγ-dependent mechanism. The increase in NPRC lowers the NPRA/NPRC ratio and reduces adipocyte natriuretic peptide signaling and results in decreased energy expenditure and impaired insulin sensitivity.

## Discussion

### Transcriptional regulation of *Nprc* in adipocytes by PPARγ via long-range distal enhancers

Although the regulation of *Nprc* expression has been linked to glucose levels, insulin action, estrogen and adrenergic receptor signaling in previous reports (29-32), our current study is the first to identify PPARγ, the master regulator of adipogenesis (19), as a transcription factor that directly controls *Nprc* expression in adipocytes. We showed that PPARγ stimulates *Nprc* expression though the action of its long-range distal enhancers rather than by direct binding at the proximal core promoter. We further demonstrated that induction of *Nprc* expression in adipose tissues by high-fat diet is also potentially mediated by PPARγ. The PPARγ enhancer RNA synthesis, an indicator of enhancer activity that could be evaluated by QPCR (28), was significantly increased in parallel to *Nprc* mRNA expression.

In our study, we compared the expression of *Nprc* with another well-characterized PPARγ target *Fabp4* (20,21). The effect of PPARγ and its agonist rosiglitazone on *Nprc* and *Fabp4* expression were not uniform in some instances. For example, under basal conditions without PPARγ agonist stimulation, the expression of *Fabp4* was significantly higher in NIH-PPARγ cells than in the parental NIH-3T3 cells, but basal *Nprc* mRNA levels were comparable in these two cell types (Fig. 2D). This suggested that *Nprc* expression is more dependent on the action of a PPARγ ligand for maximal PPARγ activation (33,34). In addition, PPARγ can bind to both the distal enhancer and the proximal promoter of *Fabp4* (21,35), which might be more potent than binding the distal enhancer alone; thus it is possible that *Fabp4* expression could still be stimulated by PPARγ without its full activation by the ligand. In line with this, the induction of *Fabp4* mRNA in NIH-PPARγ cells was also more robust than that of *Nprc* after rosiglitazone treatment (Fig. 2D). Second, while *Nprc* expression was increased in the brown fat of HFD-fed mice, *Fabp4* mRNA was actually lower even though its -5.4kb PPARγ enhancer activity was unaffected by the HFD compared to chow. This again may be due to the fact that PPARγ could bind to both *Fabp4* distal enhancer and proximal promoter (21,35), so that the activity of the enhancer alone did not fully represent its overall mRNA expression levels. Moreover, it has been reported that FABP4 can attenuate PPARγ by triggering PPARγ ubiquitination and subsequent proteasomal degradation (36). Finally, while it appears that PPARγ is the major regulator of *Fabp4* gene expression (20,21), *Nprc* gene expression is likely regulated by other factors in addition to PPARγ (29-32).

Our analysis of the long-range distal PPARγ enhancers only covered a 71kb region upstream of *Nprc* promoter. However, enhancer elements may also reside in the downstream or intronic regions. As discussed by others (37), these enhancer elements do not necessarily act on the respective closest promoter, but can bypass neighboring genes to regulate genes located more distantly along a chromosome. As such we cannot exclude the possibility that additional long-range distal elements may exist and function synergistically with the -49kb and -54kb enhancers that we identified in this study to control *Nprc* gene expression. Unbiased approaches, such as the circular chromosome conformation capture assay (38), will be required to identify these additional elements in future study.

### The physiological significance of PPARγ-dependent regulation of *Nprc* expression

Our study presents the first evidence to directly link *Nprc* gene expression to PPARγ, the master regulator of adipocyte formation. This connection may, at least partially, explain the well appreciated fact that *Nprc* is abundantly expressed in adipocytes and in the adipose tissue (6,15). It will be interesting to determine whether *Nprc* expression could also be regulated by PPARγ in the other related tissues, such as the macrophage and the cardiovascular system (39-41). In addition, we observed that *Nprc* expression was significantly higher in the male epidydimal white adipose tissue than in the female parametrial white adipose tissue, and this difference was mirrored by the PPARγ enhancer activity (data not shown). This apparent sexually dimorphic and depot-specific pattern of PPARγ activity needs further investigation.

It is well known that activation of PPARγ improves insulin sensitivity through a combination of metabolic actions, including partitioning of lipid stores and the regulation of metabolic and inflammatory mediators such as adiponectin (42). Our study showed that HFD-dependent *Nprc* expression could be potentially mediated by the action of PPARγ. An increase in NPRC expression in the adipose tissue results in reduced NPRA to NPRC ratio and thus suppresses adipose tissue natriuretic peptide signaling in obesity. As we previously showed, deletion of *Nprc* in the adipose tissue enhances NP signaling, stimulates energy expenditure, improves glucose homeostasis and protects against diet-induced obesity (6,16). Given the substantial role of NPRC for the control of natriuretic peptide signaling, our study provides a new connection to PPARγ for modulating natriuretic peptide signaling and energy metabolism during obesity.

A recently published promoter Capture Hi-C data set from human adipose tissue suggested several promoter interacting elements for the human *NPRC* gene (43). Motif and integration analysis with human adipocyte PPARγ ChIP-seq data suggested the existence of PPARγ binding sites within these NPRC promoter-interacting sequences. This indicates that *NPRC* expression may also be regulated by long-range distal PPARγ enhancers in human adipocytes. Furthermore, several studies including ours have demonstrated that adipose NPRC levels are increased with obesity in both mouse and human (9,13-18), thus the obesity-associated increase in NPRC expression is conserved between mouse and human. However, PPARγ-mediated NPRC gene expression in human adipocytes and adipose tissue will need to be further investigated.

In conclusion, our current study demonstrates that PPARγ regulates *Nprc* expression in adipocytes through its long-range distal enhancers, and the diet-dependent increase in *Nprc* expression in obesity is potentially mediated by the actions of PPARγ. Although we expect that PPARγ is not the sole regulator of *Nprc* expression in adipocytes, our study provides the first insight into the transcriptional regulation of the *Nprc* gene by establishing a new connection for how PPARγ may control natriuretic peptide signaling in the adipocyte.

## Material and Methods

### Cell lines

Mouse 3T3-L1 preadipocytes were maintained in DMEM supplemented with 10% calf serum, 100 U/ml penicillin and 100 μg/ml streptomycin. C3H10T_½_preadipocytes were maintained in DMEM supplemented with 10% fetal bovine serum (FBS), 100 U/ml penicillin and 100 μg/ml streptomycin. When reaching confluence, adipocyte differentiation was induced by a cocktail of 5µg/ml insulin, 1µM dexamethasone, 0.5mM isobutyl methylxanthine, 1µM rosiglitazone in DMEM with 10% FBS for 3 days, then maintained in DMEM with 10% FBS until day 6-7 when ready for experiment. NIH-3T3 fibroblasts and a clonal line that stably expresses PPARγ (NIH-PPARγ) were cultured as described (23,24).

### siRNA

Knockdown of *Pparγ* in differentiating 3T3-L1 adipocytes was performed with *Pparγ* esiRNA (Sigma, EMU041151) as previously described (44). Briefly, at day 4 of differentiation per the above protocol, 3T3-L1 adipocytes were trypsinized and resuspended in growth medium at a density of 2.4 × 10^6^ cells/ml. A mixture of 40 pmol siRNA and 24 μl RNAiMAX (Invitrogen) were prepared with OptiMEM medium in a final volume of 160 μl, incubated at room temperature for 5min and then added to a collagen (Gibco, A10644-01)-coated well of a 12-well plate. 3T3-L1 adipocytes were plated onto the 12-well plates at a density of 4.64×10^5^ cells with a final media volume of 0.8 ml per well for 6 hrs. After this step, cells were replaced with fresh media and cultured for another two days before harvesting for RNA and Western blot analysis.

### Chromosomal conformation capture assay (3C)

3C was performed according to previous reports (45-47). Briefly, 1 × 10^7^ C3H10T_½_adipocytes at day 6 of differentiation were trypsinized, cross-linked by 2% formaldehyde at room temperature (RT) for 10 min and quenched with 0.125 M glycine at RT for 5 min. Adipocytes were washed and lysed in a buffer (10mM Tris-HCl pH8.0, 10mM NaCl, 0.2% NP-40, 1x cOmplete Protease Inhibitor Cocktail from Roche) on ice for 30 min with shaking. The nuclei were harvested and resuspended in 0.5ml cold 1.2x NEB Buffer 2 (New England Biolabs), incubated with 0.3% SDS for 1 hr at 37°C, and with 1.8% Triton X-100 for 1 hr at 37°C. The isolated chromatin was digested with 1500 U of HindIII (NEB) overnight with shaking at 37°C and then incubated with 1.6% SDS at 65°C for 20min. The samples were transferred to 7 ml of 1.15x T4 ligation buffer (NEB), incubated with 1% Triton X-100 for 1 hour at 37°C, then ligated with 800 U T4 ligase (NEB) for 4 hours at 16°C followed by 30min at RT. The chromatin was then digested with 300 μg Proteinase K overnight at 65°C and by 30 μg RNaseA for 1 hour at 37°C. The DNA was purified with phenol/chloroform extraction, precipitated with ethanol, dissolved in 150 µl of Tris buffer (10mM, pH7.5), and quantified with NanoDrop (Thermo Scientific). Two hundred ng DNA sample was used for PCR with an anchor primer in the *Nprc* promoter fragment and a bait primer in each of the distal enhancer fragments to determine the *Nprc* promoter and enhancer interaction (Fig. 3B). The 3C PCR primers are listed in the Supplementary Table. For positive controls, BAC DNA that cover *Nprc* (RP23-305L10) and *Ercc3* (RP23-148C24) genes were digested with HindIII and randomly religated to generate all possible ligation products as described previously (47).

### Luciferase reporter assays

A -2.2kb fragment of the mouse *Nprc* gene promoter (−2233bp to +1bp, *mNprc*-2.2k) was amplified by PCR with overhangs in the forward (5’-ATGACTCGAG-3’) and reverse (5’-ATTAAAGCTT-3’) primers, digested with XhoI and HindIII and cloned into pGL-4.14 vector. The positive control plasmid with a PPARγ enhancer from *Angptl4* gene in pUC18 HSV TK-Luc vector (*Angptl4*+2.3k) was constructed as described (27,48). The individual enhancer fragments were amplified by PCR with overhangs in the forward (5’-ATTGTTGGATCC-3’) and reverse (5’-ATTGTTGTCGAC-3’) primers, digested with SalI and BamHI and cloned into pUC18 HSV TK-Luc vector. The reporter plasmid cloning primers are listed in the Supplementary Table. For luciferase assay, NIH-3T3 and NIH-PPARγ fibroblasts were plated in 12-well culture plates at a density of 1.2 × 10^5^ cells/well and reached 80% confluence the following day when ready for transfection. Cells were transfected with 1.25 μg/well of the reporter plasmids by polyethyleneimine reagent (Polysciences, #23966) and differentiated in DMEM/10% FBS with 1µM rosiglitazone for two days. Luciferase activity was assayed according to the manufacturer’s instruction (Promega, E1501) and normalized to total protein concentration (BCA, Pierce).

### High-fat diet (HFD) experiment

8-week-old wild-type male mice were fed with a high-fat diet (60% of calories from fat, Research Diet, D12492) for 12 weeks. All animal studies were approved by the Institutional Animal Care and Use Committee of Vanderbilt University Medical Center and in accordance with the NIH Guide for the Care and Use of Laboratory Animals.

### QPCR

RNA was extracted with TRIzol (Invitrogen), purified with RNeasy Mini columns (Qiagen) or Quick-RNA columns (Zymo Research) and reversed transcribed with High-Capacity cDNA Reverse Transcription Kits (Applied Biosystems). For enhancer RNA evaluation, cDNA was prepared with iScript gDNA Clear cDNA Synthesis Kit (Bio-Rad) to avoid genomic DNA contamination. QPCR was performed with PowerUp SYBR Green Master Mix (Life Technologies) according to manufacturer’s instruction. QPCR results were analyzed with the ΔΔCt method, normalized to the internal control gene 36B4 and presented as fold change relative to the control group. QPCR primers are listed in Supplementary Table.

### Western blot

Protein was extracted from cells as previously described (49). For Western blotting analysis, 30-40 µg of protein was resolved by 10% SDS–polyacrylamide gel electrophoresis, transferred to nitrocellulose membranes (Bio-Rad), incubated overnight at 4°C with specific primary antibodies in blocking buffer (Tris-Buffered Saline, 0.1% Tween-20, 5% milk or BSA) and detected with alkaline phosphatase (AP)-conjugated secondary antibody. The antibodies used were anti-NPRC (1:2000, Novus Biologicals, #NBP1-31365), anti-PPARγ (1:2000, Cell Signaling Technology, #2435), anti-FABP4 (1:1000, Cell Signaling Technology, #2120), anti-ADIPOQ (1:1000, Cell Signaling Technology, #2789), anti-GAPDH (1:2000, Protein Tech, #10494-1-AP) and anti-rabbit IgG-AP (1:20000, Sigma, #A3687)

### ChIP-seq data analysis

A previously published PPARγ ChIP-seq dataset in 3T3-L1 adipocyte and adipose tissues (Gene Expression Omnibus, GSE92606) (25) was downloaded and analyzed with Integrative Genomics Viewer (Broad Institute) (50).

### Statistical analysis

GraphPad Prism 7 was used for statistical analysis. All data were presented as means ± SEM. Unpaired two-tailed Student’s t-tests were used to determine the differences between groups. Statistical significance was defined as p-value < 0.05.

## Acknowledgements

The authors thank Ms. Wei Zhang for animal husbandry assistance, Dr. Ryan P. Ceddia for advice and comments on the manuscript, and Dr. Stephen Farmer for the NIH-3T3 and NIH-PPARγ cells.

## Author Contributions

S.C. supervised the project. F.S. designed and performed experiments and analyzed the data. Z.S. and L.N. assisted with ChIP-seq and enhancer RNA analysis. F.S. and S.C. wrote the manuscript.

## Funding and Additional Information

This work was supported by NIH R01 DK103056 (S.C.), R56 DK126890 (S.C.) and R01 DK115924 (L.N.). F.S. is supported by the America Diabetes Association Postdoctoral Fellowship Award (1-18-PDF-110). The content is solely the responsibility of the authors and does not necessarily represent the official views of the National Institutes of Health.

## Nonstandard Abbreviations

NP: natriuretic peptides,
cGMP: cyclic guanosine monophosphate,
PKG: protein kinase G,
HFD: high-fat diet,
ChIP-seq: Chromatin immunoprecipitation-sequencing,
3C: chromosomal conformation capture assay.

---

**Supplementary Table:**
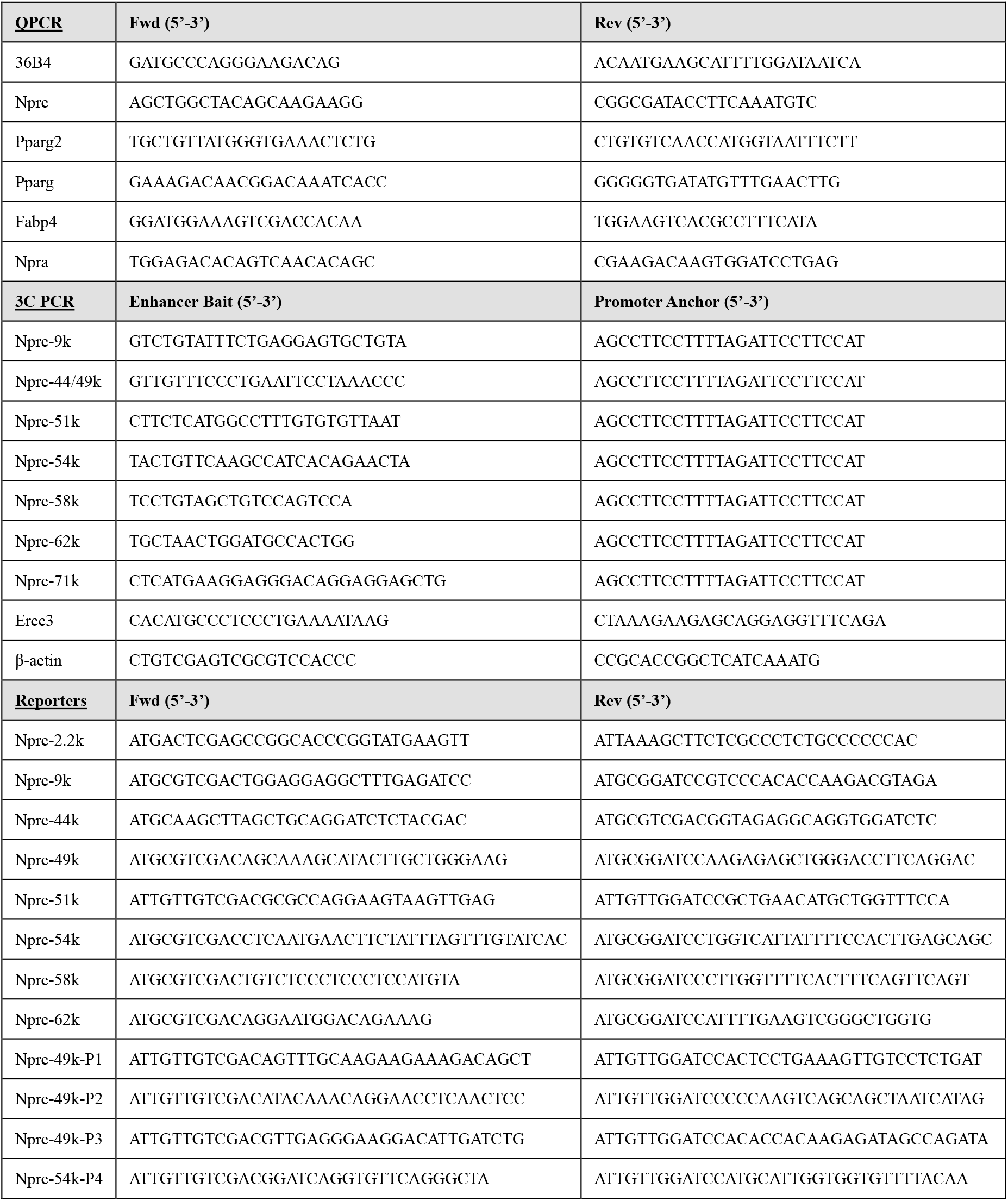
**Primer sequences**

